# A critical comparison of technologies for a plant genome sequencing project

**DOI:** 10.1101/201830

**Authors:** Pirita Paajanen, George Kettleborough, Elena López-Girona, Michael Giolai, Darren Heavens, David Baker, Ashleigh Lister, Gail Wilde, Ingo Hein, Iain Macaulay, Glenn J. Bryan, Matthew D. Clark

## Abstract

A high quality genome sequence of your model organism is an essential starting point for many studies. Old clone based methods are slow and expensive, whereas faster, cheaper short read only assemblies can be incomplete and highly fragmented, which minimises their usefulness. The last few years have seen the introduction of many new technologies for genome assembly. These new technologies and new algorithms are typically benchmarked on microbial genomes or, if they scale appropriately, human. However, plant genomes can be much more repetitive and larger than human, and plant biology makes obtaining high quality DNA free from contaminants difficult. Reflecting their challenging nature we observe that plant genome assembly statistics are typically poorer than for vertebrates. Here we compare Illumina short read, PacBio long read, 10x Genomics linked reads, Dovetail Hi-C and BioNano Genomics optical maps, singly and combined, in producing high quality long range genome assemblies of the potato species *S. verrucosum*. We benchmark the assemblies for completeness and accuracy, as well as DNA, compute requirements and sequencing costs. We expect our results will be helpful to other genome projects, and that these datasets will be used in benchmarking by assembly algorithm developers.

---

Developments in high-throughput sequencing have revolutionised genetics and genomics, with lower costs leading to an explosion in genome sequencing project size [1]. This diversity of sequencing and assembly methods, coupled to the activities of many laboratories, are generating multiple assemblies. These need to be compared to ensure that optimal approaches have been used.

The existence of very high quality references [4, 14] has made the human genome popular for demonstrating new sequencing technologies and assembly algorithms. The human genome has now been sequenced and assembled using various technologies including Sanger, 454, IonTorrent, Illumina, Pacific Biosciences (PacBio), 10x Genomics and even nanopore sequencing technologies [25, 31, 6, 37, 46, 17]. Hybrid approaches have also been used which combine complementary technologies, for example PacBio and BioNano [33].

However, the human genome is not representative of all eukaryotic genomes; plant genomes in particular are typically more repetitive (including multi-kilobase long retrotransposon elements as well as even longer regions comprising of “nested” transposon insertions). Plant biology also poses challenges for the isolation of high quality high molecular weight DNA, due to strong cell walls, co-purifying polysaccharides, and secondary metabolites which inhibit enzymes or directly damage DNA [13]. Thus technologies that work well on vertebrate genomes may not work well for plants [18]. For these reasons slow and expensive clone based minimal tiling path sequencing approaches have persisted in plants [9, 30] long after faster, cheaper short read whole genome assemblies were first demonstrated for vertebrate genomes [26]. Plant genomes also vary hugely in size, from 61 Mbp (*Genlisea tuberosa*, a member of the bladderwort family [12]) to 150 Gbp (*Paris japonica*, a relative of lilies [32]), it is still nontrivial to design a *de novo* assembly project which involves an ensemble of technologies. Each platform comes with its own input requirements, computational requirements, quality of output and, of course, labour and materials costs.

In this paper we compare several practical *de novo* assembly projects of a self-compatible, diploid Mexican wild potato species *Solanum verrucosum* using Illumina, PacBio, BioNano, 10x Genomics and Dovetail technologies. We see how plant biology poses some additional challenges for the isolation of high quality high molecular weight DNA. The genome size of about 722 Mbp is suitable for testing many different technologies whilst keeping the costs reasonable. Using the genome of *S. verrucosum* we are able to demonstrate that repeat content does limit the contiguity of the assembly by comparing the assembly to BAC sequences, and find out which technology can resolve large repeats. As its relative *S. tuberosum* has been assembled [34], we can use synteny to analyse long-range scaffolding accuracy. We find that the long-range scaffolding can cause chimeric scaffolds for some assemblies, but not others.

Our results can be used as guidance for further sequencing assembly projects and provide a basis for comparative genome studies, as each sequencing strategy and assembly method has its own biases.

## Results

The results of this study are presented in two parts. First we compare short read (Illumina) with long read (PacBio) based assemblies. In the second part we take the best performing Illumina and PacBio assemblies, and then add longer-range scaffolding data from newer technologies, namely *in vitro* Hi-C (Dovetail), optical mapping (BioNano Genomics), and read clouds (10x Genomics Chromium) technologies. Validating the assemblies for sequence and scaffolding accuracy we find strengths and weaknesses, and that methods differ hugely in their DNA, time, computational requirements and cost.

**Figure 1:**
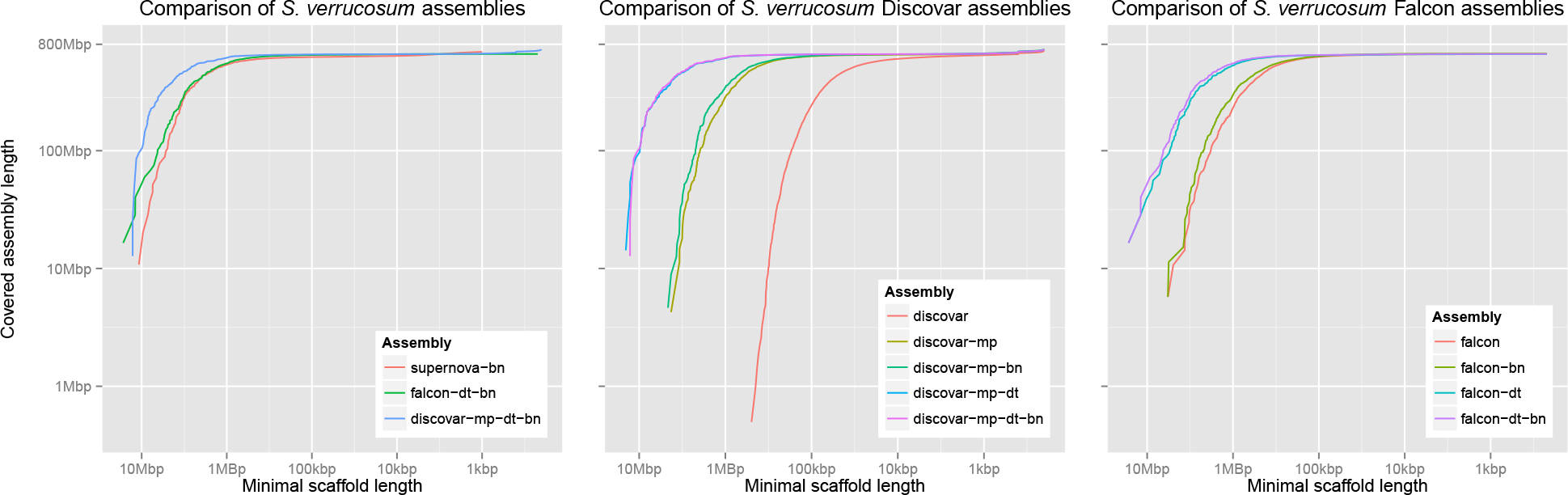
Comparison of contig/scaffold lengths and total assembly sizes of the various *S. verrucosum* assemblies

Budget constraints do play a large part in the choice of technologies to be adopted for any genome project. Assembly and scaffolding methods are often effectively the choice of sequencing method, but the properties of the genome will also affect the results. Heterozygosity, in particular, complicates the assembly process and if individual haplotypes are desired this places limitations on which strategies can be used. The careful choice of organism where possible, such as a highly inbred plant or doubled haploid, can remove the problems caused by structural heterozygosity. This approach was adopted for the potato DM reference, whereby a completely homozygous “doubled monoploid” was used instead of a highly heterozygous potato genotype. The original heterozygous diploid RH genotype selected for sequencing proved difficult to assemble due to the extremely high level of haplotype diversity.

### Contig assembly and scaffolding

The first stage of an assembly is to piece together reads to form long contiguous sequences, or *contigs* for short. These contigs can be ordered and oriented using longer-range information such as jumping/mate pair libraries. Throughout this paper we will refer to different contig assemblies that have been scaffolded. We use a naming convention which shows all of the steps used to construct the assembly. Each assembly name contains the steps used in order, separated by a hyphen. For example, the `discovar-mp-dt-bn` assembly is the `discovar` contig assembly scaffolded first with mate-pairs, then Dovetail and finally BioNano.

**Table 1:**
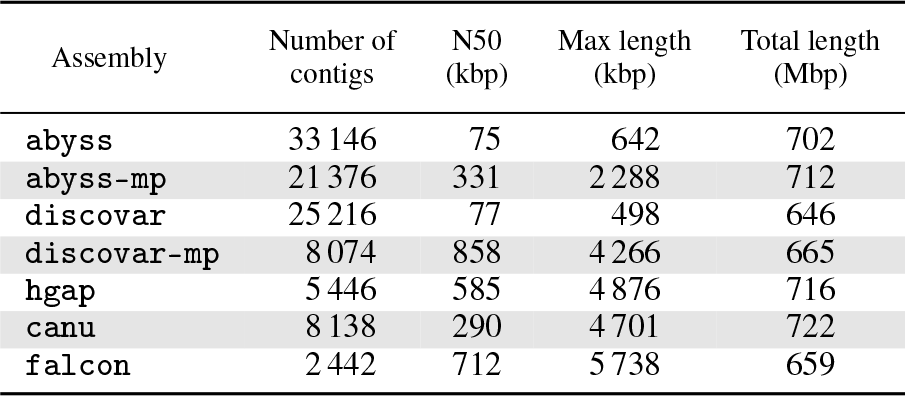
Assembly statistics of Illumina and PacBio assemblies, with a minimum contig/scaffold size of 1 kbp. `abyss` uses the TALL library, `discovar` uses the Discovar library, and `hgap, canu` and `falcon` use the PacBio library.

#### Illumina contig assembly

Two libraries were constructed for Illumina assembly. The first is a PCR-free library with insert size 500 bp (±40 %) which was sequenced with 250 bp paired-end reads on a single Illumina HiSeq run. We refer to this below as the `DISCOVAR` library. The coverage of the library was 120 ×. The second library is a PCR-free “Tight and Long Library” (TALL) with insert size 650 bp (±20 %) sequenced with 100 bp and 150 bp paired-end reads. The coverage of this library was 135 ×.

We analysed the TALL library reads with preqc, part of the SGA assembler [40], and it gave a genome size estimate at 722 Mbp, which agrees well with the 727 Mbp size of the potato genome assembly [34].

The TALL library was assembled with ABySS [41] (*k*-mer size 113) and the Discovar library using Discovar *de novo* [45] producing contig assemblies `discovar` and `abyss`, respectively. The results for these two Illumina assemblies are remarkably similar and shown in Table 1. These assemblies are more contiguous than the equivalent contig assemblies of the *S. tuberosum* genome [34].

#### Illumina scaffolding

A Nextera long mate-pair (LMP) library was made with insert size 10 000 bp (±20 %) and sequenced on two lanes of an Illumina Miseq with fragment size 500 bp and 300 bp reads. The total coverage of the LMP library was 15 ×. We scaffolded both the `discovar` and `abyss` assemblies separately using Soapdenovo2 [27] producing `discovar-mp` and `abyss-mp`, respectively. The contiguity of both was increased significantly as shown in Table 1. Here the `discovar-mp` scaffolds were slightly better so we used this assembly to take forward for longer range scaffolding with other data types.

#### PacBio assembly

A PacBio library with fragment lengths of at least 20 kbp was made giving a total coverage of 50 ×.

We conducted three long read assemblies on the same data using HGAP3 [7], part of smartanalysis (version 2.3.0p5), Canu [19] (version 1.0), and Falcon [8] (version 0.3.0) producing the `hgap, canu` and `falcon` assemblies, respectively. The assembly statistics for each is shown in Table 1.

**Figure 2:**
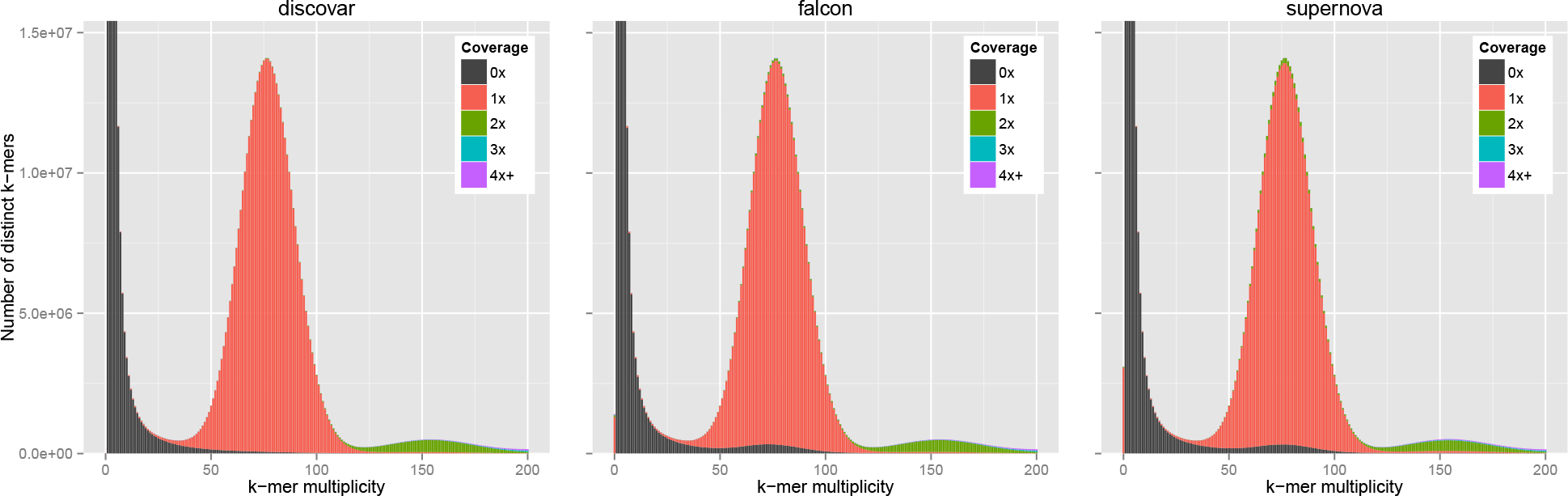
KAT spectra-cn plots comparing three *S. verrucosum* contig assemblies. The heights of the bars indicate how many *k*-mers of each multiplicity appear in the raw `DISCOVAR` reads. The colours indicate how many times those *k*-mers appear in the respective assemblies with black being zero times and red being one time. A coloured bar at zero multiplicity indicates *k*-mers appearing in the assembly which do not appear in the reads. The `FALCON` assembly has been polished with the Illumina reads using Pilon to reduce the affect of using a different sequencing platform.

The Canu assembly was made with reads that were first error-corrected by the HGAP3 pipeline because the first attempt using raw reads resulted in an excessive amounts of small scaffolds and a genome size more than 50 % longer than expected.

The `canu` and `hgap` assemblies contain considerably more content than all other assemblies. The `falcon` assembly has the highest N50, and is closest to the estimated genome length. `FALCON` also produced 9.9 Mbp of alternate contigs, likely from residual heterozygosity. We chose the `falcon` assembly to take forward to hybrid scaffolding. We first polished it using Quiver as part of SMRTanalysis (version 2.3.0p5).

### Longer-range scaffolding

To achieve higher contiguity, newer technologies have been developed to complement the previous methods and. in some cases, each other. In this section we investigate using longer range scaffolding methods to increase the contiguity of the Illumina `discovar-mp` assembly and the `falcon` PacBio assembly. We also investigate the 10x Genomics Chromium platform, an integrated solution which can be used to generate short Illumina reads with long-range positional information.

#### Dovetail

Dovetail Genomics provides a specialised library preparation method called Chicago and an assembly service using a custom scaffolder called HiRise. The Chicago library preparation technique is based on the Hi-C method, producing deliberately “chimeric” inserts linking DNA fragments from distant parts of the original molecule [35]. This is followed by standard Illumina paired-end sequencing of the inserts. Since the separation of the original fragments follows a well-modelled insert size distribution, the scaffolder is able to join contigs to form scaffolds spanning large distances, even up to 500 kbp [35].

Dovetail Genomics, LLC (Santa Cruz, CA, USA) received fresh leaf material from us from which they constructed a Chicago library. This was sequenced at Earlham Institute using Illumina 250 bp paired-end reads. The total read coverage of the Chicago library was 105×. Dovetail used their HiRise software to further scaffold the `discovar-mp` assembly, increasing the N50 from 825 kbp to 4700 kbp, and the falcon assembly, increasing the N50 from 710 kbp to 2800 kbp. These assemblies are called `discovar-mp-dt` and `falcon-dt`, respectively.

#### BioNano

The BioNano Genomics Irys platform constructs a physical map using very large DNA fragments digested at known sequence motifs with a specific nicking enzyme, to which a polymerase adds a fluorescent nucleotide. The molecules are scanned, and the distance between nicks generates a fingerprint of each molecule which is then used to build a whole genome physical map. Sequence-based scaffolds or contigs can be integrated by performing the same digestion *in silico* then ordering and orienting the contigs according to the physical map [16].

We collected BioNano data from 16 runs by repeatedly running the same chip. After filtering fragments less than 100 kbp, the yield varied from 0.8 Gb to 25.8 Gb, with the earlier runs yielding more whereas the molecule N50 was higher in later runs (ranging from 135 kbp to 240 kbp). The total yield of BioNano data was 252 Gbp which is roughly equivalent to 350× coverage.

We performed hybrid scaffolding on the `discovar-mp` and `falcon` assemblies. The *in silico* digest suggested a label density of 8.1/100 kbp for `discovar-mp` and 8.4/100 kbp for `falcon` whilst the actual observed density was only 6.8/100 kbp. We used the BioNano pipeline (v2.0) to scaffold `discovar-mp`, increasing the N50 from 825 kbp to 1260 kbp, and `falcon`, increasing the N50 from 710 kbp to 1500 kbp. These assemblies are called `discovar-mp-bn` and `falcon-bn`, respectively.

#### 10x Genomics

10x Genomics provides an integrated microfluidics based platform for generating linked reads (a cloud of non-contiguous reads with the same barcode from the same original DNA molecule) and customised software for their analysis [46]. Large fragments of genomic DNA are combined with individually barcoded gel beads into micelles in which library fragments are constructed and then sequenced as a standard Illumina library. Using the barcodes the reads from the same gel bead can be grouped together.

Unlike the previous two longer-range scaffolding approaches, the 10x Genomics platform constructs a new paired-end library which can be sequenced and then assembled into large scaffolds by one assembly program: Supernova.

**Figure 3:**
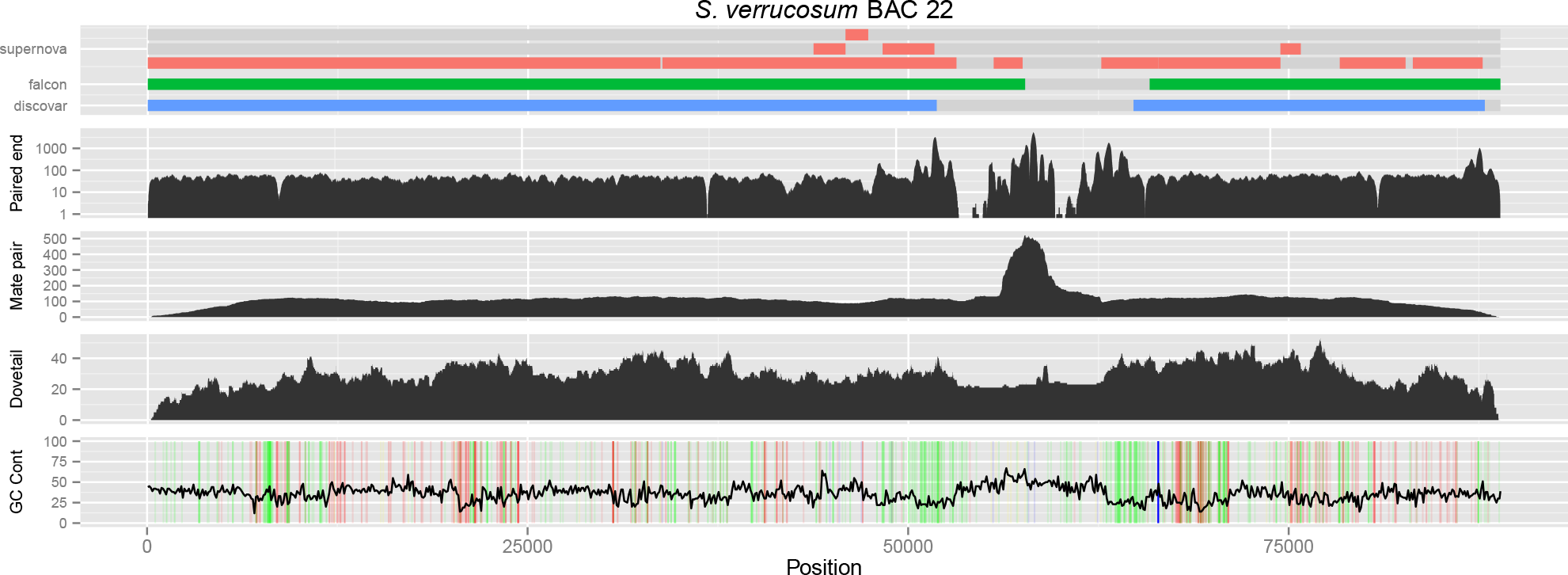
A difficult region of the genome which is contiguously assembled with a PacBio BAC but in none of our whole genome assemblies. The region was correctly scaffolded by Dovetail. The figure shows various alignments and information with respect to the BAC assembly. The top track shows the contigs which appear in the `discovar`, `falcon` and `supernova` assemblies. The paired-end track shows read coverage of the `DISCOVAR` paired-end library. The mate-pair and Dovetail tracks show physical/fragment coverage of the mate-pair and Dovetail libraries, respectively. The bottom track shows GC content of the sequence as well as homopolymers sequences of at least 5 bp where A, C, G, and T are coloured as red, blue, yellow, and green, respectively.

A 10x Genomics Chromium library was made according to manufacturer’s instructions and a lane of Illumina HiSeq 250 bp paired-end reads were generated with a coverage of about 92 ×. Supernova (version 1.1.1) produced the `supernova` assembly with length 641 Mbp and a scaffold N50 of 2.33 Mbp. Trimming reads back to 150 bp or reducing sequencing depth to 56 ×, which are the read length and depth recommended by 10x Genomics, generated very similar results (see Supplemental Section 2.3).

#### Hybrid scaffolding

It is possible to iteratively combine these longer-range scaffolding approaches. We tested several hybrid approaches using the `discovar`, `falcon` and `supernova` assemblies. For example the `discovar-mp` assembly was scaffolded using Dovetail and then BioNano producing `discovar-mp-dt-bn` with an N50 of 7.0 Mbp, the highest contiguity of any assembly reported here. The `falcon` assembly when scaffolded with both produced scaffolds with an N50 of only 3.09 Mbp, lower than with BioNano alone. Finally we scaffolded the `supernova` assembly with BioNano producing `supernova-bn` which increased the N50 from 2.33 Mbp to 2.85 Mbp.

We also used long reads from PacBio to scaffold and to perform “gapfilling” on the assemblies, replacing regions of unknown sequence (N stretches) with a PacBio consensus sequence. This also presents an opportunity to use lower coverage PacBio data to improve an Illumina assembly, which may be more cost effective than a *de novo* assembly using PacBio. PBJelly (version 15.2.20) [11] was used to perform gapfilling using only 10 SMRTcells of PacBio data (8 × depth). The `Supernova` assembly increased in size from 641 Mbp to 671 Mbp, and N50 from 2.33 Mbp to 2.64 Mbp, and the amount of Ns present reduced from 7.58 % to 5.14%. The `discovar-mp-dt` assembly increased in size from 656 Mbp to 680 Mbp and N50 from 4.69 Mbp to 4.87 Mbp, with Ns reduced from 3.03 % to 1.28 %. However, how gaps and percentage Ns are generated differs between assembly methods (see
Discussion).

## Assembly evaluation

Achieving a genome assembly with high levels of contiguity is potentially useless if it does not faithfully represent the original genome sequence. We assessed errors in assemblies by comparison to the raw data used to make the assemblies, as well as measuring gene content, local accuracy (BAC assemblies), and long-range synteny with the close relative *Solanum tuberosum.*

### K-mer content

Analysis of the *k*-mer content of an assembly gives a broad overview of how well the assembly represents the underlying genome. We used the PCR-free Illumina Discovar library as our reference for the *k*-mer content of the genome. Due to the high accuracy of the reads we expect the *k*-mer spectra for a library to form a number of distributions which correspond to read errors, non-repetitive, and repetitive content in the genome. These distributions can be seen by observing only the shapes and ignoring the colours in Figure 2. The reader is referred to the KAT documentation for further details [29].

In Figure 2 we compare the *k*-mer contents of the three contig assemblies—`discovar`, `falcon`, and `supernova`—to the `DISCOVAR` library. To minimise the effects of the differences between Illumina and PacBio sequencing error profiles the `falcon` assembly has been polished with the Illumina reads using Pilon [44] (see Supplemental Figure S3.1 for the unpolished plot).

The small red bar on the origin in some plots shows content which appears in the assembly but not in the Illumina reads. The `discovar` assembly is very faithful to the content in the library. The black area denotes sequences in the reads but not in the assembly: those clustering at the origin are predicted sequence errors in the reads, the small amount between 50–100 on the *x*-axis is sequence missing from the assembly. The dominant red peak (1 ×, around multiplicity 77), which is the vast majority of all assemblies here, contains content in the Illumina reads which appears once in the assembly (homozygous sample). Green areas on top of the main peak in `FALCON` and `SUPERNOVA` represents possible duplications in the assembly, whereas the green (2 ×) small peak to the right of the main peak is probably true duplicates— as these sequences are present twice in the assembly and at twice the expected read counts. At the main peak (*k*-mer multiplicity 77), the amount of potentially duplicated content in the assemblis is 0.66 % in `falcon`, 1.3 % in `supernova`, and 0.15 % in `discovar`.

### Gene content

We assessed the gene content of the three most contiguous assemblies—`discovar-mp-dt-bn`, `falcon-dt-bn`, and `supernova-bn`—using two datasets. The first is with `BUSCO` and its `embryophyta_odb9` (plants) dataset [39] and the second is all the predicted transcript sequences from the *S. tuberosum* genome [34].

We found that each of the three assemblies shows at least 95 % of `BUSCO`s as complete, with only 2–3 % missing. The difference is small but the `discovar-mp-dt-bn` assembly is the most complete while `supernova-bn` is the worst performing. The results are shown in Figure 4.

We aligned the *S. tuberosum* representative transcript sequences to each genome assembly using `BLAST` [2] and then measured how much of each transcript sequence was represented in the assembly according to various minimum percentage identity cutoffs. As expected when comparing between species, as the threshold approaches 100% nucleotide identity the transcript completeness drops closer to zero. Using a threshold between 96–98 % we find the median transcript completeness is highest in `discovar-mp-dt-bn`, followed by `falcon-dt-bn`, and then `supernova-bn`. However, the difference between the assemblies is small, Figure 5 shows a box and whisker plot of completeness of the representative transcript sequences.

### Local accuracy

As BACs are easier to assemble due to smaller size and a much more limited amount of repetitive DNA content than a whole genome, we assessed the performance of our three assemblies at a local scale using BAC assemblies. We randomly selected, sequenced, and assembled 96 BAC clones from *S. verrucosum* BAC library. We chose 20 high-quality BAC assemblies (single scaffolds/contigs with Illumina or PacBio) to measure the accuracy of the whole genome assemblies.

We used dnadiff [20] to compare the BAC sequences to the `supernova-bn`, `discovar-mp-dt-bn`, and `falcon-dt-bn` assemblies finding sequence identities of 99.40 %, 99.97 %, and 99.87 %, respectively. As in the previous section, the `discovar-mp-dt-bn` assembly shows the highest accuracy, with `supernova-bn` the lowest, though the differences are small.

To illustrate the performance of the different technologies sequencing different genomic features we mapped whole genome reads and assemblies to single BACs as shown in Figure 3. None of our three whole genome assemblies are able to reconstruct BAC 22; each breaking at a large (more than 12kbp) repeat. The Discovar library (paired-end), mate-pair library and Dovetail library were each mapped and only reads mapping to a high quality and exhibiting up to one mismatch are shown in the figure. The mapping reveals several areas of high repetition, for example the arms and middle of a retrotransposon, and there are areas lacking coverage completely which suggests a sequence which is difficult for our Illumina sequence data to resolve. We also see drops in coverage at some sites with high concentrations of homopolymers, as marked by coloured lines in the GC content, for example an A rich region of ~7 kbp. Interestingly the repeat arms are also rich in homopolymers.

**Figure 4:**
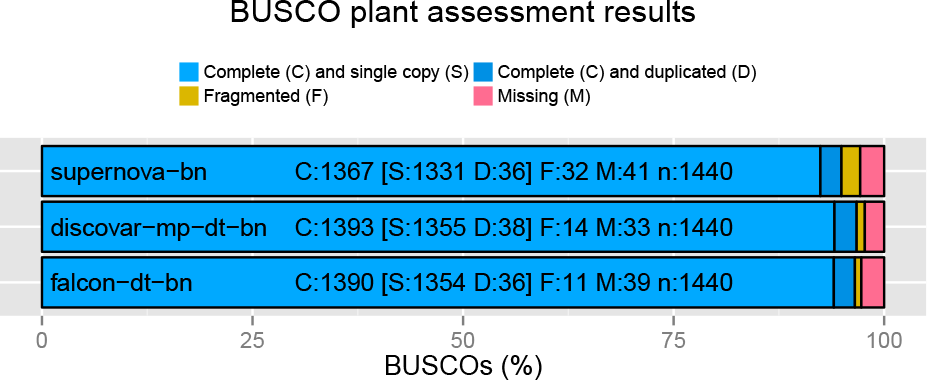
Busco analysis of `supernova-bn`, `discovar-mp-dt-bn`, and `falcon-dt-bn` using the plant gene dataset.

**Figure 5:**
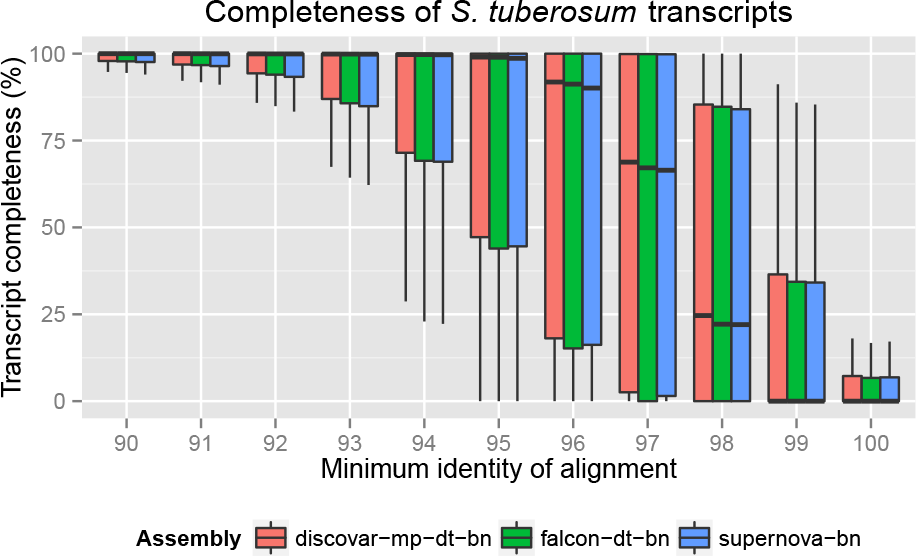
Box and whisker plot showing completeness of the *S. tuberosum* transcripts in `supernova-bn`, `discovar-mp-dt-bn`, and `falcon-dt-bn` with various levels of minimum percentage identity.

We note that the `discovar-mp-dt-bn` assembly leaves the largest gap around the repeat. The `falcon` assembly was able to completely cover an area with no mapping paired-end Illumina reads which explains some of extra *k*-mer content in Figure 2 noted earlier in this assembly. The `supernova-bn` assembly was able to reconstruct more of the difficult region, but it also contains duplications in the homopolymer rich flanking regions that is not seen in the other assemblies.

The mate-pair library was not able to scaffold the `discovar` contigs due to the size of this repeat being larger than its 10 kbp insert size. The mate-pair fragments also map to a great depth in the repeat. Dovetail data, however, shows a much smoother fragment distribution and was able to scaffold the two `discovar` contigs in the correct order and orientation as it could scaffold up to 50 kbp (the cutoff used by the HiRise scaffolder). However, the gap length was not estimated with Dovetail and was arbitrarily set to 100 Ns when in reality the gap is over 12 000 bp long.

### Long-range accuracy using synteny to *S. tuberosum*

As all our assemblies are *de novo,* in the sense that we used no prior information from other Solanaceae genomes, we reasoned that more accurate long range scaffolding would be apparent as longer syntenic blocks to a closely related species. We used nucmer [20] to analyse the synteny of our assemblies to the pseudomolecules of the *S. tuberosum* genome [38]. Figure 6 shows the mummer plot for chromosome 11 of *S. tuberosum* against our three assemblies. We saw the `falcon-dt-bn` assembly showed the best synteny with the `discovar-mp-dt-bn` being the worst. The plots for the remaining chromosomes are shown in Supplemental Figures S3.2, S3.3, and S3.4.

**Figure 6:**
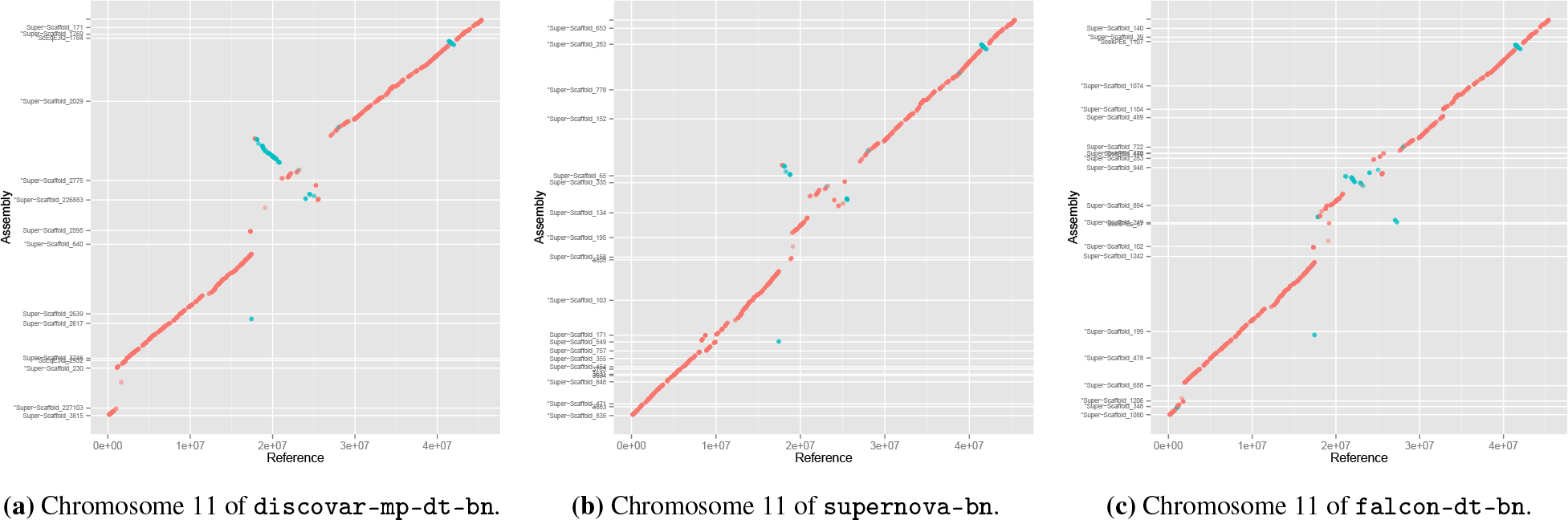
Mummer plots showing alignment to chromosome 11 of the *S. tuberosum* reference. The *S. tuberosum* reference is shown on the *x*-axis and assembly scaffolds on the *y*-axis. Alignments shown are at least 10 kbp long and 90% identical.

Using synteny we identified two cases of chimerism, i.e. scaffolds that align well to two different pseudomolecules of *S. tuberosum* genome. Both cases are in `discovar-mp-dt-bn` but not `falcon-dt-bn`. The first 1.5 Mbp of scaffold ScEqE3Q_528 maps to pseudomolecule 7 while the last 2.9 Mbp map to pseudomolecule 2 in the *S. tuberosum* genome. There is no conflict reported with the BioNano Genomics optical map in this area, but we can exclude the possibility that these are real chromosome structural arrangements in *S. verruscosum* because we have GbS markers on each end of this scaffold which also map in an *S. verrucosum* cross to these different linkage groups (López-Girona unpublished). The other case is a scaffold ScEqE3Q_633 in which the first 1.4 Mbp map to pseudomolecule 8 and the remainder to pseudomolecule 3, here BioNano Genomics does report a conflict which would highlight this error, and *S. verrucosum* genetic markers also support the chimera classification.

## Discussion

A Discovar assembly is the cheapest and easiest to construct, and the resulting assembly is very accurate, albeit highly fragmented. Adding a long mate-pair library is a proven method of increasing the contiguity of a short read assembly by scaffolding. The 10x Genomics based assembly using Supernova was as easy to obtain as the Discovar assembly. The two most remarkable features of this assembly are the low cost and input DNA requirement: for only slightly higher cost than a Discovar assembly, and considerably less than with only one long mate-pair library, we obtained an assembly comparable to what one would expect from multiple long mate-pair libraries.

Our PacBio assembly using Falcon achieved contiguity similar to that of `discovar-mp` (Discovar plus long mate-pair scaffolding). PacBio sequencing has a considerably higher cost and material requirement than Illumina sequencing, but the Falcon assembly contains truly contiguous sequence as opposed to `discovar-mp` which contains gaps patched with Ns. The PacBio read lengths (N50=13.5kbp) were similar to the insert size of mp library (mean 10 kb), and the read coverage was higher for PacBio (50 ×) than for the mp data (15 x), but PacBio contigs (N50=712kbp) are slightly shorter than the `discovar-mp` scaffolds (N50=858 kbp).

The addition of Dovetail showed the most striking increase in contiguity by scaffolding. We note that our Dovetail scaffolds provided the order and orientation of the constituent contigs but no estimate for the length of the gaps between them. This should be taken into consideration if true physical length of sequences is important, and for specific downstream uses. Both Illumina (Discovar+Mp) and PacBio (`FALCON`) assemblies are amenable to the addition of Dovetail, but the scaffolds produced from the Falconcontigs (4 × increase) were not as long as those from the Illumina assembly (5.5 × increase). This could be because while the Falconassembly has been polished with PacBio reads, it retains some PacBio errors and so some Dovetail (Illumina) reads do not pass stringent mapping filters. If true, Pilon polishing with Illumina reads could help, as it improved the *k*-mer spectra (Figure 2).

With BioNano Genomics restriction enzyme digest based optical maps we obtained less (~2 × increase) scaffolding improvement than with Dovetail (4–5.5 × increase). This could be due to three issues: first that assembly gaps are not correctly sized which prevents real, and *in silico,* restriction maps matching (as information is purely encoded in the distances between sites). We see that the ungapped PacBio assemblies improve more than scaffolded Illumina, and Dovetail scaffolds (with arbitrary 100 bp gaps) hardly increase at all. Secondly, because the method produces low information density (one enzyme site per ~12kbp) long fragments with many sites are need to create significant matches, and our DNA was not sufficiently long (best run N50 was 240 kbp). Longer DNA (over 300 kbp), and perhaps multiple enzyme maps with iterative scaffolding could have improved the results. Thirdly we observe that the *in silico* restriction rates for Illumina and PacBio assemblies are similar (8.1–8.4 sites /100kbp) whereas the actual observed rates from the physical map is much lower at 6.8 sites/100 kbp, suggesting that there could be a fraction of the genome missing from our assemblies which is very low in sites such as centromeric or telomeric regions where the BioNano Genomics map can not scaffold through.

Gapfilling using PBJelly offers an attractive method of using the long read data from PacBio to improve an existing Illumina based assembly. This closed many of the gaps in the scaffolds thereby decreasing the fraction of unknown sequence (Ns) and also increasing the contiguity. The increase in contiguity of the 10x Genomics assembly was the highest. It will be intriguing to see if an assembly approach combining Chromium data with long reads (directly on the assembly graph) can combine the best attributes of both data types to resolve complex regions.

Analysis of the *k*-mer content of the `supernova`, `discovar`, and `falcon` assemblies showed that the *k*-mer spectra of each assembly is very clean. We see slightly higher level of sequence duplication in the `supernova` assembly, and to a lesser extent in the `falcon` assembly. All three assembly algorithms are diploid aware, meaning they are able to preserve both haplotypes. The gene content of each assembly was very similar with all three of our long assemblies showing a high percentage of the expected genes. The 10x Genomics based assembly showed a slightly lower count in both of our assessments but the difference is very small.

We used multiple BAC assemblies of ~100kb insert size to illustrate the technical limitations of each method. Short read methods cannot resolve many areas of repetition within a WGS assembly. This is especially noticeable in a plant genome with higher repeat content, and is one of the major reasons for breaks in contiguity in these assemblies. In our example in Figure 3, the long mate-pair library alone is not sufficient. It takes the larger fragment lengths within the Dovetail Chicago library to finally make the join in the whole genome assembly.

Long read technologies do not suffer as much with repeats and, in the case of PacBio, tend to have more random rather than systematic errors [5]. We can see in our examplar that the `falcon` assembly covers some of the repetitive region. The underlying BAC assembly was also obtained with PacBio and gave us a single true contig for the entire BAC. On close inspection we noticed that difficult region was spanned by reads of length 22–26 kbp. This shows that long reads are certainly able to span such regions of difficulty, and to assemble them. Recently ultra-long reads with an N50 of 99.7 kbp (max. 882kbp) with ~92% accuracy have been produced with the new MinION R9.4 chemistry using high molecular weight DNA [17]. If this is also achievable on plant material the remaining repetitive fraction of genomes should become visible.

To evaluate the longer range accuracy of our genome assemblies we compared them to the closely related *S. tuberosum* pseudomolecule assembly, which revealed good synteny with all three of our longest assemblies (`discovar-mp-dt-bn`, `falcon-dt-bn` and `supernova`). There are some disagreements especially in the centromeric areas, but as these appeared in all assemblies these could illustrate real structural variation. We detected two chimeric scaffolds in the `discovar-mp-dt-bn` assembly but neither is present in the `falcon-dt-bn`. The two Dovetail scaffolding processes shared the same Hi-C sequence data but were conducted many months apart (`discovar-mp` first and later `falcon`), so may use different versions of Dovetail’s proprietary HiRise software. On detailed examination we see that the ScEqE3Q_528 scaffold chimeric join is made by Dovetail hopping through a fragmented area of short (1–2 kbp) contigs. Such small contigs do not exist in the `Falcon` assembly, which maybe why we do not find chimeras. BioNano Genomics finds it hard to map to areas with many Dovetail gaps (as these are set to an arbitrary 100 bp size), and this region also has a high enzyme nicking rate (nearly twice the genome average), including two areas where nicks are less than 200 bp apart and so would be optically merged. In scaffold ScEqE3Q_633 we detect that `discovar-mp` scaffold123 was correctly split by Dovetail data as chimeric (also highlighted by BioNano Genomics and genetic markers) but the scaffold was not broken at the exact chimeric join, and the remaining sequence from the wrong chromosome was sufficient for Dovetail to propagate the error. Whilst we did not detect a high level of systematic errors in any of our assembly methods, the importance of using BioNano Genomics and genetic markers to identify chimeras that then can be broken is apparent.

**Table 2:**
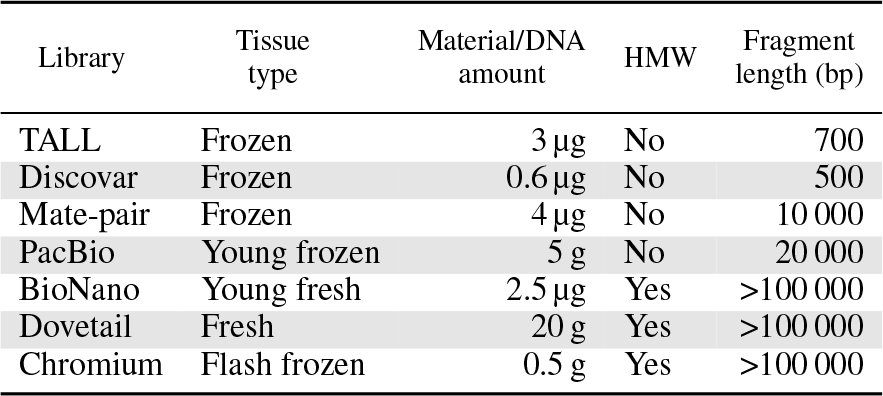
Material requirements for each library. Amounts in grams are for fresh/frozen material and amounts in micrograms for DNA. In each case where frozen or flash frozen is stated, fresh material is also acceptable.

## Materials and Methods

### Project requirements

Each of the assembly methods we have used comes with its own requirements. We have broken this down into material requirements, that is plant and DNA material, monetary requirements, that is the cost of preparation and sequencing, and computational requirements. Table 2 lists the material requirements for each library.

We calculated costs taking into consideration the costs of consumables, laboratory time, and machine overheads, but not bioinformatics time. For sequencing costs we used the Duke University cost as much as possible to provide comparative figures. Since several of the projects share common methods, such as sequencing a lane on a HiSeq 2500, we have broken down the costs into individual components. See Table 3 for our full costs calculations.

In many cases the assemblies can be performed with modest scientific computing facilities. In some cases, notably for Supernova, a very large amount of memory is required. In this case the computing requirement will not be available to most laboratories and will need to be sourced elsewhere. Table 4 shows the computational requirements of each assembly method.

### Library preparation and sequencing

In this section we briefly describe methods for library preparation and sequencing. For a comprehensive description, please see the supplementary material.

*S. verrucosum* accesssion Ver-54 was grown in the glass house in James Hutton Institute in Scotland. Both fresh and frozen leaves from this accession and its clones were used for DNA extraction.

The TALL library was prepared using 3 μg of DNA and fragments of 650 bp were sequenced with a HiSeq2500 with a 2 × 150 bp read metric. The Discovar library was prepared using 600 ng of DNA and fragments of 500 bp were sequenced with a HiSeq2500 with a 2 × 250 bp read metric.

**Table 3:**
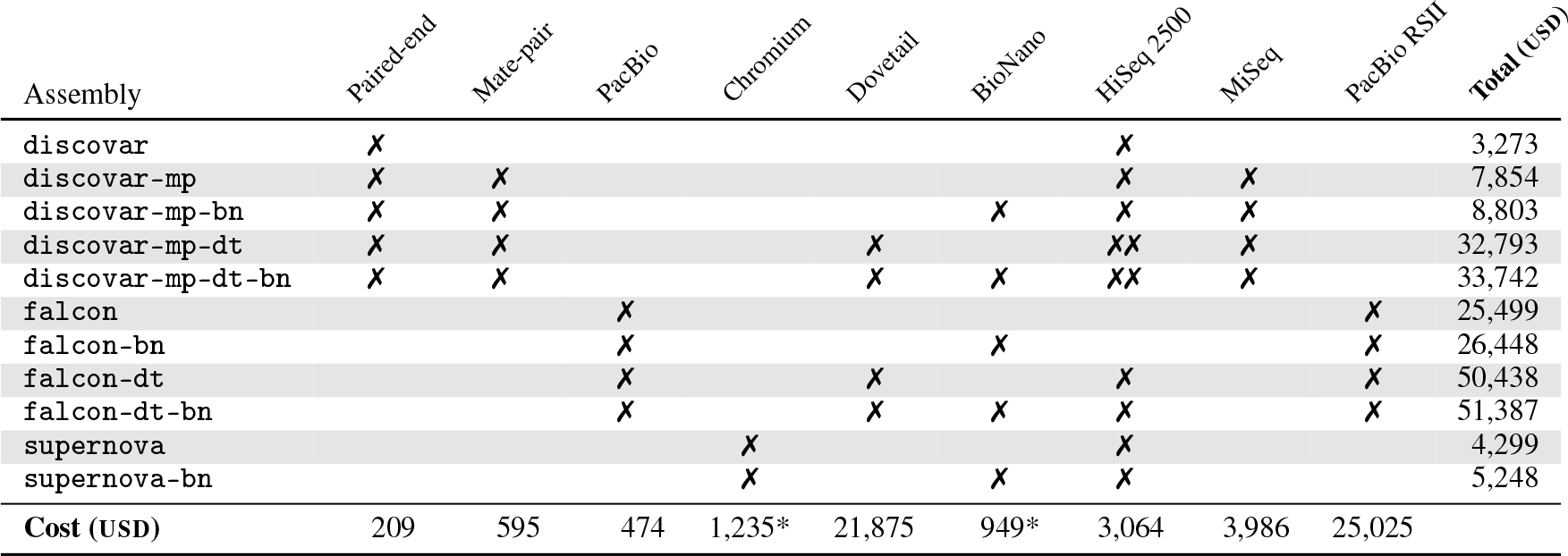
The overall cost of each assembly project. We show which library preparations and sequencing runs are required for each assembly with a checkmark (✗). Individual costs are given at the bottom, and total costs of each assembly on the right. All costs are according to Duke University as of April 2017 and in USD, except those marked with a * which were according to the Earlham Institute and converted from GBP to USD at an exchange rate of 0.804 GBP/USD. Paired-end, mate-pair, PacBio, and Chromium are library preparations including DNA extraction. Dovetail includes Chicago library preparation and HiRise scaffolding. BioNano is the cost of building the optical map. HiSeq2500 is for a rapid run half flowcell (one lane) with 250 bp reads. MiSeq is for two runs with 300 bp reads. PacBio RSII is for 65 SMRT cells.

**Table 4:**
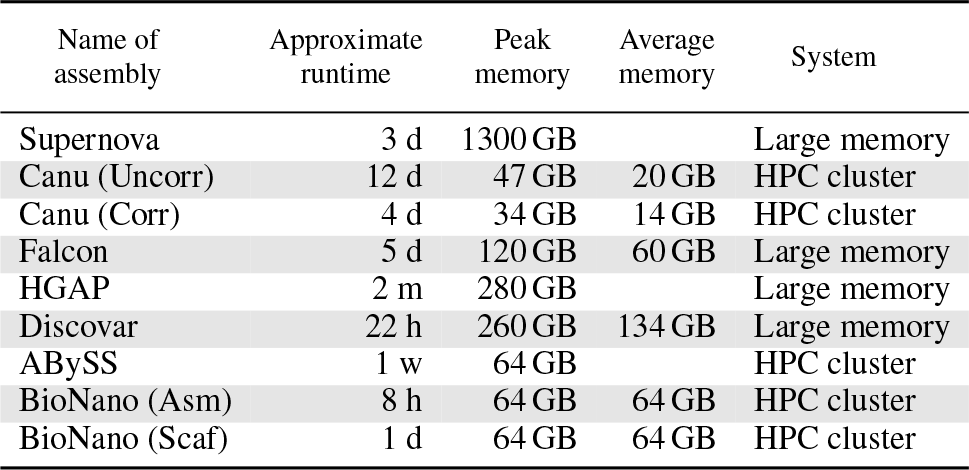
Computational requirements.

The mate-pair library was prepared using 4 μg of DNA and fragments of 10 kbp were circularised, fragmented and sequenced on a MiSeq with a 2×300 bp read metric.

A PacBio library was prepared using 5 g of frozen leaf material. A 20 kbp fragment length library was prepared according to manufacturer’s instructions and sequenced on 65 SMRT cells with the P6C4 chemistry on a PacBio RSII.

The 10x Chromium library was prepared according to the manufacturer’s instructions and sequenced on a HiSeq2500 with a 2×250 bp read metric.

For BioNano, DNA was extracted using the IrysPrep protocol. 300 ng was used in the Nick, Label, Repair and Stain reaction and loaded onto a single flow cell on a BioNano chip. The chip was run eight times to generate 252 Gb of raw data.

### Assembly and evaluation

All tools and scripts that were used to perform the evaluation and produce the figures are available on GitHub in the georgek/potato-figures repository.

We used RAMPART [28] to run `ABySS` [41] multiple times with different *k* values. Discovar *de novo* was run with normal parameters.

Long mate-pair reads were first processed with NextClip [22] to remove the Nextera adapter. Soapdenovo2 was then used to perform scaffolding with both the paired-end and mate-pair libraries. *k*-mer content was analysed with the `kat comp` tool [29]. We used default parameters with manually adjusted plot axes to show the relevant information.

We used the Busco core plant dataset to evaluate the gene content. The *S. tuberosum* representative transcripts were aligned to the assemblies using `BLAST` and the coverage of transcripts at various thresholds using a tool we developed.

The BACs were sequenced with the Earlham Institute BAC pipeline [3] and were assembled with Discovar *de novo* using normal parameters after filtering for *E. coli* and the BAC vector. The PacBio BAC was assembled using HGAP. We used GNU parallel [42] for concurrent assembly and analysis.

20 BACs which assembled into a single contig were selected to use as a reference. These BACs are non-redundant to the extent that they do not share any lengths of sequence of more than 95 % identity and over 5000 bp long. Short reads were aligned to the BACs using Bowtie2 [21] with default parameters. The assemblies were mapped to the BACs using `bwa mem` [23]. The mapped sequences were sorted and filtered for quality using sambamba [43]. Fragment coverage was calculated using samtools [24] and bedtools [36].

Synteny was analysed with mummer [10]. We used nucmer to align the assemblies to the *S. tuberosum* reference v4.04 [15]. Alignments less than 10 kbp and 90 % identity were filtered out.

## Data Access

All read data generated in this study have been submitted to the EMBL-EBI European Nucleotide Archive under the project PRJEB20860.

## Acknowledgements

We thank Lawrence Percival-Alwyn and Walter Verweij for their assistance in library preparation and analysis, and Michael Bevan for critical reading of this manuscript. This work was funded with BBSRC project grants (BB/K019325/1) and (BB/K019090/1). This work was strategically funded by the BBSRC, Core Strategic Programme Grant (BB/CSP17270/1) at the Earlham Institute. High-throughput sequencing and library construction was delivered via the BBSRC National Capability in Genomics (BB/CCG1720/1) at the Earlham Institute (EI, formerly The Genome Analysis Centre, Norwich), by members of the Platforms and Pipelines Group. This research was supported in part by the NBI Computing infrastructure for Science (CiS) group through the HPC cluster and UV systems. We thank Duke University for providing sequencing costs via Dugsim (https://dugsim.net/).

## Authors’ contributions

GB, ELG, IH, and GW prepared the sample. MDC, GK, and PP designed the analysis. DB, GB, ELG, MG, DH, IH, AL, and IM constructed libraries and performed sequencing. GK and PP made the assemblies and GK, ELG, and PP performed the evaluation. MDC, GK, GB, ELG and PP wrote and prepared the manuscript. All authors read and approved the final manuscript.

## References

1. The 1000 Genomes Project Consortium. “An integrated map of genetic variation from 1,092 human genomes”. In: Nature 491.7422 (2012), pp. 56–65. ISSN: 0028-0836. doi: 10.1038/nature11632.

2. S. F. Altschul et al. “Basic local alignment search tool”. In: Journal of Molecular Biology 215.3 (1990), pp. 403–410. ISSN: 0022-2836. DOI: 10.1016/S0022-2836(05)80360-2.

3. S. Beier et al. “Construction of a map-based reference genome sequence for barley, Hordeum vulgare L.” In: Scientific Data 4 (2017). DOI: 10.1038/sdata.2017.44.

4. E. Callaway. “‘Platinum’ genome takes on disease”. In: Nature News 515.7527 (2014), p. 323. DOI: 10.1038/515323a.

5. M. O. Carneiro et al. “Pacific biosciences sequencing technology for genotyping and variation discovery in human data”. In: BMC Genomics 13.1 (2012), p. 375. ISSN: 1471-2164. DOI: 10.1186/1471-2164-13-375..

6. M. J. P. Chaisson et al. “Resolving the complexity of the human genome using single-molecule sequencing”. In: Nature 517.7536 (2015), pp. 608–611. ISSN: 0028-0836. doi: 10.1038/nature13907.

7. C.-S. Chin et al. “Nonhybrid, finished microbial genome assemblies from long-read SMRT sequencing data”. In: Nature Methods 10.6 (2013), pp. 563–569. ISSN: 1548-7091. doi: 10.1038/nmeth.2474.

8. C.-S. Chin et al. “Phased diploid genome assembly with singlemolecule real-time sequencing”. In: Nature Methods 13.12 (2016), pp. 1050–1054. ISSN: 1548-7091. DOI: 10.1038/nmeth.4035.

9. F. Choulet et al. “Structural and functional partitioning of bread wheat chromosome 3B”. In: Science 345.6194 (2014), pp. 1249721–1249721. ISSN: 0036-8075. doi: 10.1126/science.1249721.

10. A. L. Delcher, S. L. Salzberg and A. M. Phillippy. “Using MUMmer to identify similar regions in large sequence sets”. In: Current Protocols in Bioinformatics (2003), pp. 10–3. DOI: 10.1002/0471250953.bi1003s00.

11. A. C. English et al. “Mind the gap: upgrading genomes with Pacific Biosciences RS long-read sequencing technology”. In: PLOS ONE 7.11 (2012), e47768. ISSN: 1932-6203. doi: 10.1371/journal.pone.0047768.

12. A. Fleischmann et al. “Evolution of genome size and chromosome number in the carnivorous plant genus Genlisea (Lentibulariaceae), with a new estimate of the minimum genome size in angiosperms”. In: Annals of Botany 114.8 (2014), pp. 1651–1663. ISSN: 0305–7364. DOI: 10.1093/aob/mcu189.

13. E. A. Friar. “Isolation of DNA from plants with large amounts of secondary metabolites”. In: Methods in Enzymology. Molecular Evolution: Producing the Biochemical Data 395 (2005), pp. 1–12. ISSN: 0076-6879. DOI: 10.1016/S0076-6879(05)95001-5.

14. “Genome in a bottle—a human DNA standard”. In: Nature Biotech 33.7 (2015), pp. 675–675. ISSN: 1087-0156. doi: 10.1038/nbt0715-675a.

15. M. A. Hardigan et al. “Genome reduction uncovers a large dispensable genome and adaptive role for copy number variation in asexually propagated Solanum tuberosum”. In: The Plant Cell (2016), TPC2015–00538-RA. ISSN:, 1532-298X. doi: 10.1105/tpc.15.00538.

16. A. R. Hastie et al. “Rapid genome mapping in nanochannel arrays for highly complete and accurate de novo sequence assembly of the complex Aegilops tauschii genome”. In: PLOS ONE 8.2 (2013), e55864. ISSN: 1932-6203. doi: 10.1371/journal.pone.0055864.

17. M. Jain et al. “Nanopore sequencing and assembly of a human genome with ultra-long reads”. In: bioRxiv (2017). doi: 10.1101/128835. eprint: https://www.biorxiv.org/content/early/2017/04/20/128835.full.pdf.

18. W.-B. Jiao and K. Schneeberger. “The impact of third generation genomic technologies on plant genome assembly”. In: Current Opinion in Plant Biology 36 (2017), pp. 64–70. ISSN: 1369-5266. DOI: 10.1016/j.pbi.2017.02.002.

19. S. Koren et al. “Canu: scalable and accurate long-read assembly via adaptive k-mer weighting and repeat separation”. In: Genome Research 27.5 (2017), pp. 722–736. ISSN: 1088-9051, 1549-5469. DOI: 10.1101/gr.215087.116.

20. S. Kurtz et al. “Versatile and open software for comparing large genomes”. In: Genome Biology 5.2 (2004), R12. DOI: 10.1186/gb-2004-5-2-r12.

21. B. Langmead and S. L. Salzberg. “Fast gapped-read alignment with Bowtie 2”. In: Nature Methods 9.4 (2012), pp. 357–359. doi: 10.1038/nmeth.1923.

22. R. M. Leggett et al. “Nextclip: an analysis and read preparation tool for Nextera long mate pair libraries”. In: Bioinformatics 30.4 (2014), pp. 566–568. ISSN: 1367-4803. doi: 10. 1093/bioinformatics/btt702.

23. H. Li. “Aligning sequence reads, clone sequences and assembly contigs with BWA-MEM”. In: arXiv preprint arXiv:1303.3997 (2013).

24. H. Li et al. “The sequence alignment/map format and SAMtools”. In: Bioinformatics 25.16 (2009), pp. 2078–2079. DOI: 10.1093/bioinformatics/btp352.

25. R. Li et al. “De novo assembly of human genomes with massively parallel short read sequencing”. In: Genome Research 20.2 (2010), pp. 265–272. ISSN: 1088-9051, 1549-5469. doi: 10.1101/gr.097261.109.

26. R. Li et al. “The sequence and de novo assembly of the giant panda genome”. In: Nature 463.7279 (2010), pp. 311–317. ISSN: 0028-0836. DOI: 10.1038/nature08696.

27. R. Luo et al. “SOAPdenovo2: an empirically improved memory-efficient short-read de novo assembler”. In: GigaScience 1 (2012), p. 18. ISSN: 2047-217X. DOI: 10.1186/2047-217X-1-18.

28. D. Mapleson, N. Drous and D. Swarbreck. “Rampart: a workflow management system for de novo genome assembly”. In: Bioinformatics 31.11 (2015), pp. 1824–1826. ISSN: 1367-4803. doi: 10.1093/bioinformatics/btv056.

29. D. Mapleson et al. “KAT: a k-mer analysis toolkit to quality control NGS datasets and genome assemblies”. In: Bioinformatics (2016). DOI: 10.1093/bioinformatics/btw663.

30. M. Mascher et al. “A chromosome conformation capture ordered sequence of the barley genome”. In: Nature 544.7651 (2017), pp. 427–433. ISSN: 0028-0836. DOI: 10.1038/nature22043.

31. Y. Mostovoy et al. “A hybrid approach for de novo human genome sequence assembly and phasing”. In: Nature Methods 13.7 (2016), pp. 587–590. ISSN: 1548-7091. DOI: 10.1038/nmeth.3865.

32. J. Pellicer, M. F. Fay and I. J. Leitch. “The largest eukaryotic genome of them all?” In: Botanical Journal of the Linnean Society 164.1 (2010), pp. 10–15. ISSN: 0024-4074. doi: 10.1111/j.1095-8339.2010.01072.x.

33. xgiv Pendleton et al. “Assembly and diploid architecture of an individual human genome via single-molecule technologies”. In: Nature Methods 12.8 (2015), pp. 780–786. ISSN: 1548-7091. doi: 10.1038/nmeth.3454.

34. The Potato Genome Sequencing Consortium. “Genome sequence and analysis of the tuber crop potato”. In: Nature 475.7355 (2011), pp. 189–195. ISSN: 0028-0836. DOI: 10.1038/nature10158.

35. N. H. Putnam et al. “Chromosome-scale shotgun assembly using an in vitro method for long-range linkage”. In: Genome Research 26.3 (2016), pp. 342–350. ISSN: 1088-9051, 1549-5469. DOI: 10.1101/gr.193474.115.

36. A. R. Quinlan and I. M. Hall. “BEDTools: a flexible suite of utilities for comparing genomic features”. In: Bioinformatics 26.6 (2010), pp. 841–842. DOI: 10.1093/bioinformatics/btq033.

37. J. M. Rothberg et al. “An integrated semiconductor device enabling non-optical genome sequencing”. In: Nature 475.7356 (2011), pp. 348–352. ISSN: 0028-0836. DOI: 10.1038/nature10242.

38. S. K. Sharma et al. “Construction of reference chromosome-scale pseudomolecules for potato: integrating the potato genome with genetic and physical maps”. In: G3&#58; GeneslGenomeslGenetics 3.11 (2013), pp. 2031–2047. ISSN: 2160-1836. DOI: 10.1534/g3.113.007153.

39. F. A. Simão et al. “BUSCO: assessing genome assembly and annotation completeness with single-copy orthologs”. In: Bioinformatics 31.19 (2015), pp. 3210–3212. ISSN: 1367-4803. doi: 10.1093/bioinformatics/btv351.

40. J. T. Simpson and R. Durbin. “Efficient de novo assembly of large genomes using compressed data structures”. In: Genome Research 22.3 (2012), pp. 549–556. DOI: 10.1101/gr.126953.111.

41. J. T. Simpson et al. “ABySS: a parallel assembler for short read sequence data”. In: Genome Research 19.6 (2009), pp. 1117–1123. DOI: 10.1101/gr.089532.108.

42. O. Tange. “GNU parallel—the command-line power tool”. In: ;login: The USENIX Magazine 36.1 (2011), pp. 42–47. DOI: 10.5281/zenodo.16303.

43. A. Tarasov et al. “Sambamba: fast processing of NGS alignment formats”. In: Bioinformatics 31.12 (2015), pp. 2032–2034. doi: 10.1093/bioinformatics/btv098.

44. B. J. Walker et al. “Pilon: an integrated tool for comprehensive microbial variant detection and genome assembly improvement”. In: PLOSONE 9.11 (2014), pp. 1–14. DOI: 10.1371/journal.pone.0112963.

45. N. I. Weisenfeld et al. “Comprehensive variation discovery in single human genomes”. In: Nature Genetics 46.12 (2014), pp. 1350–1355. ISSN: 1061-4036. DOI: 10.1038/ng.3121.

46. N. I. Weisenfeld et al. “Direct determination of diploid genome sequences”. In: Genome Research (2017). ISSN: 1088-9051, 1549-5469. DOI: 10.1101/gr.214874.116.

